# The influence of environmental setting on the community ecology of Ediacaran organisms

**DOI:** 10.1101/861906

**Authors:** Emily G. Mitchell, Nikolai Bobkov, Natalia Bykova, Alavya Dhungana, Anton Kolesnikov, Ian R. P. Hogarth, Alexander G. Liu, Tom M.R. Mustill, Nikita Sozonov, Shuhai Xiao, Dmitriy V. Grazhdankin

## Abstract

The broad-scale environment plays a substantial role in shaping modern marine ecosystems, but the degree to which palaeocommunities were influenced by their environment is unclear. To investigate how broad-scale environment influenced the community ecology of early animal ecosystems we employed spatial point process analyses to examine the community structure of seven bedding-plane assemblages of late Ediacaran age (558–550 Ma), drawn from a range of environmental settings and global localities. The studied palaeocommunities exhibit marked differences in the response of their component taxa to sub-metre-scale habitat heterogeneities on the seafloor. Shallow-marine palaeocommunities were heavily influenced by local habitat heterogeneities, in contrast to their deep-water counterparts. Lower species richness in deep-water Ediacaran assemblages compared to shallow-water counterparts across the studied time-interval could have been driven by this environmental patchiness, because habitat heterogeneities correspond to higher diversity in modern marine environments. The presence of grazers and detritivores within shallow-water communities may have promoted local patchiness, potentially initiating a chain of increasing heterogeneity of benthic communities from shallow to deep-marine depositional environments. Our results provide quantitative support for the “Savannah” hypothesis for early animal diversification – whereby Ediacaran diversification was driven by patchiness in the local benthic environment.

**Author Contributions:** E. Mitchell conceived this paper and wrote the first draft. N. Bobkov, A. Kolesnikov, N. Sozonov and D. Grazhdankin collected the data for DS surface. N. Bobkov and N. Sozonov performed the analyses on DS surface. N. Bykova, S. Xiao, and D. Grazhdankin collected the data for WS, KH1 and KH2 surfaces and E. Mitchell performed the analyses. A. Dhungana and A. Liu collected the data for FUN4 and FUN5 surfaces and A. Dhungana performed the analyses. T. Mustill and D. Grazhdankin collected the data for KS and T. Mustill and E. Mitchell performed the analyses. I. Hogarth developed the software for preliminary KS surface analyses. E. Mitchell, N. Bobkov, N. Bykova, A. Dhungana, A. Kolesnikov, A. Liu, S. Xiao and D. Grazhdankin discussed the results and prepared the manuscript.

## Background

The Ediacaran–Cambrian transition (∼580–520 million years ago) is one of the most remarkable intervals in the history of life on Earth, witnessing the rise of large, complex animals in the global oceans (1, 2). The diversification of early animals coincides with dramatic perturbations in the global abiotic environment, including changes to carbon cycling and a progressive but dynamic oxygenation of the oceans (3, 4). The extent to which animals themselves drove these global changes is a matter of considerable debate (5–7) with several competing hypotheses suggested to explain their observed diversification. These include global abiotic changes that occured over kilometre scales (8, 9) and biotic factors acting over local scales (metre to kilometre), and include organism interactions such as burrowing and/or predation (10, 11). Feedbacks between biotic and abiotic factors have also been proposed as drivers of early animal diversification, whereby Ediacaran organisms directly or indirectly created patchy food resources, stimulating the evolution of mobile bilaterians (12, 13). Due to the small (within community) spatial scales over which key evolutionary mechanisms often act (14), investigation of the community ecology of Ediacaran assemblages over broad (kilometre) spatial scales offers an opportunity to link the interactions of individual organisms to macro-evolutionary and macro-ecological trends. In this study, we investigate the relationship between late Ediacaran early animal diversification and the broad-scale environment.

Ediacaran macrofossils occur globally across a wide-range of palaeo-environments (1). Previous studies have separated late Ediacaran palaeocommunities into three taxonomically distinct assemblages – the Avalon, White Sea and Nama – which occupy partially overlapping temporal intervals and different water-depths with no significant litho-taphonomic or biogeographic influence (15–17). This study focusses on palaeocommunities within the Avalon and White Sea fossil assemblages that are considered to reflect original in situ communities (18, 19), permitting the use of statistical analyses of the distribution of fossil specimens on bedding planes (spatial point process analyses, SPPA) to reconstruct the interaction of organisms with each other and their local environment (20–25). The Avalon assemblage is primarily represented by sites in Newfoundland, Canada and Charnwood Forest UK (26, 27), and typically documents mid-shelf/deep-water settings (from depths below the edge of the continental shelf – the slope break) of 575–566 Ma (28, 29). Such sites exhibit relatively limited ecological and morphological diversity (30, 31), and palaeocommunities consisting almost exclusively of sessile taxa (32) that show only weak trends with community composition along regional palaeoenvironment gradients (20). Previous spatial analyses of Avalonian communities have found limited evidence for environmental interactions within these communities (21–23), in contrast to the strong imprint exerted by resource-limitation on modern deep-sea ecosystems (33, 34).

Palaeocommunities from the White Sea assemblage are most famously represented by sites in South Australia, and the East European Platform of Russia, dating to ∼558–555 Ma (35–37). These assemblages typically document shallow-water, diverse communities including taxa interpreted as bilaterians (38), herbivores (39), detritivores (40) and motile organisms (41). Within the White Sea assemblages, community composition is strongly correlated with sedimentary environment and the presence of textured organic surfaces at bed-scale level (42, 43).

Metrics of taxonomic and ecological diversity are much higher in White Sea assemblages than in Avalonian ones, with changes in taxonomic and morphological diversity calculated to be of similar magnitude to those between the Ediacaran and Cambrian (30, 31). These Ediacaran assemblages have high beta-diversity compared to modern benthic systems (44), but the driving processes underlying this high diversity are not understood. The regional palaeoenvironment (kilometre scale) (15, 17) has a significant influence on (non-algal dominated) Ediacaran fossil assemblage composition, but metreits influence on local (metre to sub-metre scale) community ecology has not yet been investigated. In modern benthic communities, small spatial scale (< 50 cm) substrate heterogeneities (e.g. substrate variations in nutrients, oxygen patchiness, or biotic and abiotic gradients within microbial mats) exert a significant influence on community ecology (33,34,45). For Ediacaran palaeocommunities, it is not possible from spatial analyses alone to determine the underlying causes of habitat heterogeneities, nor the extent to which they relate to food resources, such as those resulting from the decay of Ediacaran organisms (12, 46). However, it is possible to compare how the relative influence of such heterogeneities changes with broad-scale environmental setting: previous analyses have identified assemblage-level trends between community compositions and bathymetric depth (15–17). In this study, we compare the drivers of community ecology between shallow and deep-water Ediacaran palaeocommunities (above or below the slope break) over a ∼7-million-year period using spatial analyses of seven palaeocommunities.

### Spatial analyses

Determining the nature of interactions between fossilised organisms and their environment can be undertaken if entire palaeocommunities are preserved in-situ, such that the position of the fossils on bedding planes can be interpreted to reflect aspects of the organism’s life-history (47). For sessile organisms, such as in the Avalon communities, community-scale spatial distributions are dependent upon the interplay of a limited number of factors: physical environment (which manifests as habitat associations of a taxon or taxon-pairs (48)); organism dispersal/reproduction (49); competition for resources (50); facilitation between taxa (where one taxon increases the survival another taxa) (51); and differential mortality (52). For fossil assemblages containing mobile taxa (e.g. the White Sea assemblages), behavioural ecology also influences spatial distributions, so interpretations of their spatial distributions are qualitative rather than quantitative.

Studies of modern ecosystems have demonstrated that habitat associations resulting from interactions between organisms and their local environment can be either positive, leading to aggregations of individuals (such as around a preferential substrate for establishment), or negative segregation away from such patches (21). SPPA are a suite of analyses compare the relative density of points (in this case fossil specimens) to different models corresponding to different ecological processes, in order to infer the most likely underlying process responsible for producing the observed spatial distribution. For sessile organisms, habitat associations identified by SPPA are best-modelled by a heterogeneous Poisson model (HP), or when combined with dispersal limitations, an Inhomogeneous Thomas Cluster model (ITC) (53, 54). Where the local environment is resource-limited to the extent that it significantly reduces organism densities, this is indicated by spatial segregation between specimens within a community (55). When sessile populations are not significantly affected by their local environment, their spatial distributions are completely spatially random (CSR), indicating no significant influence by any biological or ecological processes at the spatial scale investigated, or alternatively reflect dispersal/reproductive processes (48,54,56–58). CSR is modelled by homogeneous Poisson processes (47), whereas dispersal patterns are best modelled by best-fit Thomas Cluster (TC) or Double Thomas Cluster (DTC) models (54). Facilitation (where one taxa increases the survival of another) is best-modelled by linked-cluster models (51, 59) and density-dependent processes detected using random-labeling analyses (52, 60).

### Geological setting

We assessed the community palaeoecology of seven fossil-bearing assemblages across five different global Ediacaran locations, spanning the full range of known habitats inhabited by members of the Ediacaran macrobiota during the late Ediacaran interval, and incorporated data from previous studies (21, 23) on Avalonian palaeocommunities for comparison. These localities document a range of diverse local depositional environments, but in order to focus on the broadest macro-ecological and macro-evolutionary patterns we have coarsely grouped them within either shallow or deep-water settings.

#### Shallow marine settings

Five of the studied palaeocommunities are found in facies that reflect shallow marine depositional environments. Palaeocommunity WS is an *Aspidella*–bearing surface on the underside of a wave-rippled sandstone within a thick package of mudstones and sandstones deposited in a prograding, storm-influenced depositional system (61, 62). It was collected from the Lyamtsa Formation of the Valdai Group, along the Onega Coast of the White Sea, Russian Federation, and remained in the field where it was destroyed by landslides. *Aspidella* specimens were collected and are stored uncatalogued at the Trofimuk Institute for Petroleum Geology and Geophysics in Novosibirsk. The Lyamtsa Formation is older than a date of 558 ± 1 Ma (U/Pb zircon dating of volcanic tuffs near the base of the overlying Verkhovka Formation) (16). Surface (KS) is on the lower surface of a finely laminated sandstone, interpreted as a flood deposit within a prograding prodelta depositional system (63). This surface, within the lower member of the Erga Formation (Winter Coast of the White Sea) (16, 35), contains the fossil *Kimberella*, and is younger than 552.85 ± 0.77 Ma (64) (date recalculated from Martin et al. (65)). The KS surface remained in the field and has been subsequently destroyed by land slides and weathering. Two *Funisia*-bearing surfaces from the base of thin-bedded wave-rippled quartz sandstones representing deposition in prodelta marine settings between fair-weather and storm wave base originate from the Ediacara Member of South Australia (42,66–68). These surfaces reside in the collections of the South Australia Museum, with surface FUN4 collected from Ediacara Conservation Park (SAM P55236) and surface FUN5 collected from the Mount Scott Range (SAM P41506). Since FUN4 and FUN5 originate from different localities (> 50 km apart), is it assumed likely that they represent discrete bedding plane/palaeocommunities. The South Australian Ediacaran successions have not been radiometrically dated, but the Ediacara Member is widely assumed to be of a similar age to the White Sea fossil-bearing sections (1, 2).

Surface DS is a *Dickinsonia*-bearing surface from the Konovalovka Member of the Cherny Kamen Formation, cropping out along the Sylvitsa River, Central Urals, Russia (63, 69). It lies within an interval of finely alternating wave-rippled sandstones, siltstones and mudstones that are sandwiched between two thick intervals of biolaminated sandstone characterised by microbial shrinkage cracks and salt crystal pseudomorphs (70). The overall succession is considered transitional from marginal marine to non-marine, with the fossil-bearing interval interpreted as having been deposited in a lagoon within a tidal flat depositional system (70). A U/Pb zircon date of 557 ± 13 Ma from volcanic tuffs near the base of the Cherny Kamen Formation (63) suggests that this unit may have been deposited broadly coevally with those on the White Sea coast. Specimens from this surface reside in Novosibirsk State University, Russian Federation (specimen numbers: 2057-001 to 2057-003) and will be placed at the Ural Geological Museum (Yekaterinburg).

All five of these surfaces therefore represent siliciclastic depositional environments from above the slope break, and so fall broadly into the grouping of “shallow marine”. They contain examples of taxa interpreted as animals (e.g. *Dickinsonia*, *Kimberella*) as well as non-metazoans (*Orbisiana*) and their age and facies place them within the White Sea assemblage (15, 17).

#### Deep-water marine setting

Two bedding surfaces dominated by *Aspidella* specimens (KH1 and KH2) were collected from a package of finely alternating limestone and shale interbeds within the Khatyspyt Formation, Olenek River, Siberia. Sedimentological observations (e.g., turbiditic nature of the limestones; evidence of strong unidirectional flows; intraclasts originating from outside of the Khatyspyt depositional basin) suggest the Khatyspyt Formation was deposited within a starved intracratonic rift basin developed in a marine ramp setting within a relatively deep-water setting beyond the shelf slope break (71–74). A positive δ^13^C_carb_ excursion in the Khatyspyt Formation has been correlated with an excursion of similar magnitude in the <550 Ma Gaojiashan Member of the Dengying Formation (74). Strontium isotope ratios (^87^Sr/^86^Sr) in the Khatyspyt Formation are consistently ca. 0.7080 (74, 75), a value approaching some of the ratios seen in the Gaojiashan Member (76), so this correlation seems plausible. Surface KH2 remains in the field and surface KH1 was destroyed while excavated KH. Specimens from KH1 surface reside in Trofimuk Institute for Petroleum Geology and Geophysics, collection number 913 (specimen numbers: 0607/2009-3, 0607/2009-6, 0607/2009-7, 0607/2009-17, 0607/2009-18).

### Data Collection

Spatial data were collected from the surfaces using different methods depending on the physical properties of the bedding plane. The WS, KH1, KH2 surfaces were mapped in the field (WS in 2017, KH1 in 2006 and 2009, and KH2 in 2018) onto millimetre graph paper. First, the co-ordinates of the edge of the rock surface were recorded, then the co-ordinates, orientation and dimensions of each of the specimen were measured and plotted onto the paper. For DS, a bedding surface of 9 m^2^ was excavated over the course of two years (2017–2018). The surface was photo-mapped, with photographs taken under an artificial light source at night. The intersection between maximum length (L) and maximum width (W) of each specimen was taken to be the absolute position of the organism, with measurements obtained from digital photographs using Adobe Photoshop CC software and Apple Script Editor.

The KS surface was excavated in July 2004, and is a laterally discontinuous transect consisting of four slabs of variable size, ranging from 0.6 × 0.4 m to 1.6 × 1.0 m. The relative positions of the slabs within the transect were mapped in situ on an excavated terrace. A separate block originating from the same horizon was found in float close to the transect. Following reassembly, the taxonomic identity, positions, orientations and shapes of the fossils were mapped at millimetre scale. For the FUN4 and FUN5 surfaces, photogrammetric maps of the bedding surfaces were made, with lens edge effects corrected using RawTherapee (v. 2.4.1). For all mapped palaeocommunities, fossil identification, position, and dimensions (disc width, disc length, stem length, stem width, frond length, and frond width) were digitized in Inkscape 0.92.3 on a 2D projection of the dataset, resulting in a 2D vector map for each palaeocommunity. Only taxa that had sufficient abundance (> 5 specimens) for spatial analyses were formally identified, and these were grouped within one of six taxonomic groups: *Aspidella*, *Dickinsonia*, *Funisia*, *Kimberella*, *Orbisiana,* and the trace fossil *Kimberichnus*. A group consisting of all the sessile taxa on the KS surface was also assessed, because abundance was not sufficient to include all taxa individually. Analyses were not conducted for individual low abundance taxa whose specimen numbers fell below the threshold for which results would be statistically meaningful.

## Methods

### Bias analyses

For each surface, we first tested for erosional biases and tectonic deformation, since both have the potential to distort spatial analyses (18, 73). If these factors were found to have significantly affected specimen density distributions, the erosion and/or deformation were taken into account when performing later analyses (cf. (23)), with heavily eroded sections of the bedding planes excluded from analyses. The influence of tectonic deformation was only observed on the DS surface, so retrodeformation techniques (18, 25) were not applied to the spatial maps of WS, KH1, KH2, KS, FUN4 and FUN5 surfaces. Where possible (WS, KH1 and KH2 surfaces), the area near the outcrops was investigated, and no independent evidence for tectonic deformation was found. The holdfast discs on surfaces KS, FUN4 and FUN5 did not show any evidence tectonic deformation. The DS surface showed signs of deformation in the form of consistent variation in specimen length to width ratios along a presumed axis of deformation. The *`fitModel`* function from the *`mosaic`* package in R (73) was used to find the best-fit values for the direction and strength of deformation using the assumption that *Dickinsonia* had a consistent length to width ratio during the ontogeny (43,77,78) though note (79)), and the spatial map was retrodeformed cf. (18,23,25).

### Spatial Analyses

Initial data exploration, inhomogeneous Poisson modelling, and segregation tests were performed in R (75) using the package *`spatstat`* (*80, 81*). Programita was used to obtain distance measurements and to perform aggregation model fitting (described in detail in references (*48, 52, 80, 82–86*).

Univariate and bivariate pair correlation functions (PCFs) were calculated from assemblage population densities using a grid of 1 cm × 1 cm cells on all surfaces except DS, where a 10 cm × 10 cm cell size was used to correspond to the larger overall mapped area. To minimise noise, a 3 cell smoothing was calculated dependent on specimen abundance, which was applied to the PCF (59). To test whether the PCF exhibited complete spatial randomness (CSR), 999 simulations were run for each univariate and bivariate distribution, with the 49 highest and 49^th^ lowest values removed (59). CSR was modelled by a Poisson model on a homogeneous background where the PCF = 1 and the fit of the fossil data to CSR was assessed using Diggle’s goodness-of-fit test (56, 87). Note that due to non-independence of spatial data, Monte-Carlo generated simulation envelopes cannot be interpreted as confidence intervals. If the observed data fell below the Monte-Carlo simulations, the bivariate distribution was interpreted to be segregated; above the Monte-Carlo simulations, the bivariate distribution was interpreted to be aggregated (47, 59).

If a taxon was not randomly distributed on a homogeneous background, and was aggregated, the random model on a heterogeneous background was tested by creating a heterogeneous background from the density map of the taxon under consideration. This density map was defined by a circle of radius R over which the density was averaged throughout the sample area. Density maps were formed using estimators within the range of 0.1 m < *R* < 1 m, with *R* corresponding to the best-fit model used. If excursions outside the simulation envelopes for both homogeneous and heterogeneous Poisson models remained, then Thomas cluster models were fitted to the data as follows:

1. The PCF and L-function (88) of the observed data were found. Both measures were calculated to ensure that the best-fit model is not optimized towards only one distance measure, and thus encapsulates all spatial characteristics.
2. Best-fit Thomas cluster processes (89) were fitted to the two functions where PCF > 1. The best-fit lines were not fitted to fluctuations around the random line of PCF = 1 in order to aid good fit about the actual aggregations, and to limit fitting of the model about random fluctuations. Programita used the minimal contrast method (56, 87) to find the best-fit model.
3. If the model did not describe the observed data well, the lines were re-fitted using just the PCF. If that fit was also poor, then only the L-function was used.
4. 99 simulations of this model were generated to create simulation envelopes, and the fit checked using the O-ring statistic (82).
5. In order to assess how well the model fit the observed data, the goodness-of-fit (*p_d_*) was calculated over the model range (86). A *p_d_ = 0* indicates no model fit, and *p_d_ = 1* indicates a perfect model fit. Very small-scale segregations (of the order of specimen diameter) were not included in the model fitting, since they likely represent the finite size of the specimens, and a lack of specimen overlap.
6. If there were no excursions outside the simulation envelope and the *p_d_*-value was high, then a univariate homogeneous Thomas cluster model was interpreted as the best model.

For any univariate distributions exhibiting CSR, the size-classes of each taxon were calculated, the univariate PCFs of the smallest size-classes and largest size-classes were plotted, with 999 Monte Carlo simulations of a complete spatially random distribution and segregation tests performed. The most objective way to resolve the number and range of size classes in a population is by fitting height-frequency distribution data to various models, followed by comparison of (logarithmically scaled) Bayesian information criterion (BIC) values (86), which we performed in R using the package `MCLUST` (90). The number of populations identified was then used to define the most appropriate size classes. A BIC value difference of >10 corresponds to a “decisive” rejection of the hypothesis that two models are the same, whereas values <6 indicate only weakly rejected similarity of the models (90–94). Once defined, the PCFs for each size class were calculated.

Bivariate analyses were performed on the KS surface (the only surface with multiple abundant taxa/taxon groups) between *Kimberella* – *Orbisiana, Kimberella* – *Kimberichnus* and *Orbisiana* – *Kimberichnus*. For each taxon pair, the bivariate PCF was calculated, and then compared to CSR using Monte Carlo simulations and Diggle’s goodness-of-fit test.

## Results

Across the seven palaeocommunities, *Dickinsonia* on the DS surface was the only taxon that exhibited CSR. There were five univariate distributions (Sessile Taxa on KS, *Funisia* on FUN4 and FUN5, *Aspidella* on KH1 and KH2) exhibiting aggregated spatial distributions, two univariate (*Aspidella* on WS and large *Dickinsonia* on DS) and one bivariate (*Kimberella* and *Kimberichnus* on KS) segregated spatial distributions (Fig. 3, Table 2). The *Aspidella* aggregations from KH1 and KH2 were best modelled by the same double Thomas cluster process (*p_d_ ^kh1^ = 0.883*, *p_d_ ^k21^* = 0.932, Fig. 3G, H; Table 2), which consisted of large clusters of 20.96 cm diameter containing smaller clusters with a mean of six specimens within a cluster of 7.34 cm in diameter (Fig. 3G and H, (95)). These results indicate that the non-random spatial distributions were most likely due to two generations of reproduction cf. (47), and do not represent a significant interaction or association with local habitat variations. This result is consistent with previous work on older (∼565 Ma) deep-water communities that also show a strong non-environmentally influenced signal (23). In contrast, the *Aspidella* from the WS surface show significant segregation and are best-modelled by a heterogeneous Poisson process (*p_d_ ^ws^ = 0.796,* Fig 3F, Table 2). This is consistent with small-scale intra-specific competition in a resource-limited environment (55). *Funisia* from FUN4 and FUN5 had aggregations that are best-modelled by heterogeneous Poisson processes (*p_d_ ^Fun4^ =0.9570*, *p_d_ ^Fun5^ = 0.9080*, Fig. 3 D, E; Table 2), which are interpreted to indicate significant habitat associations with the local environment.

**Fig. 1.**
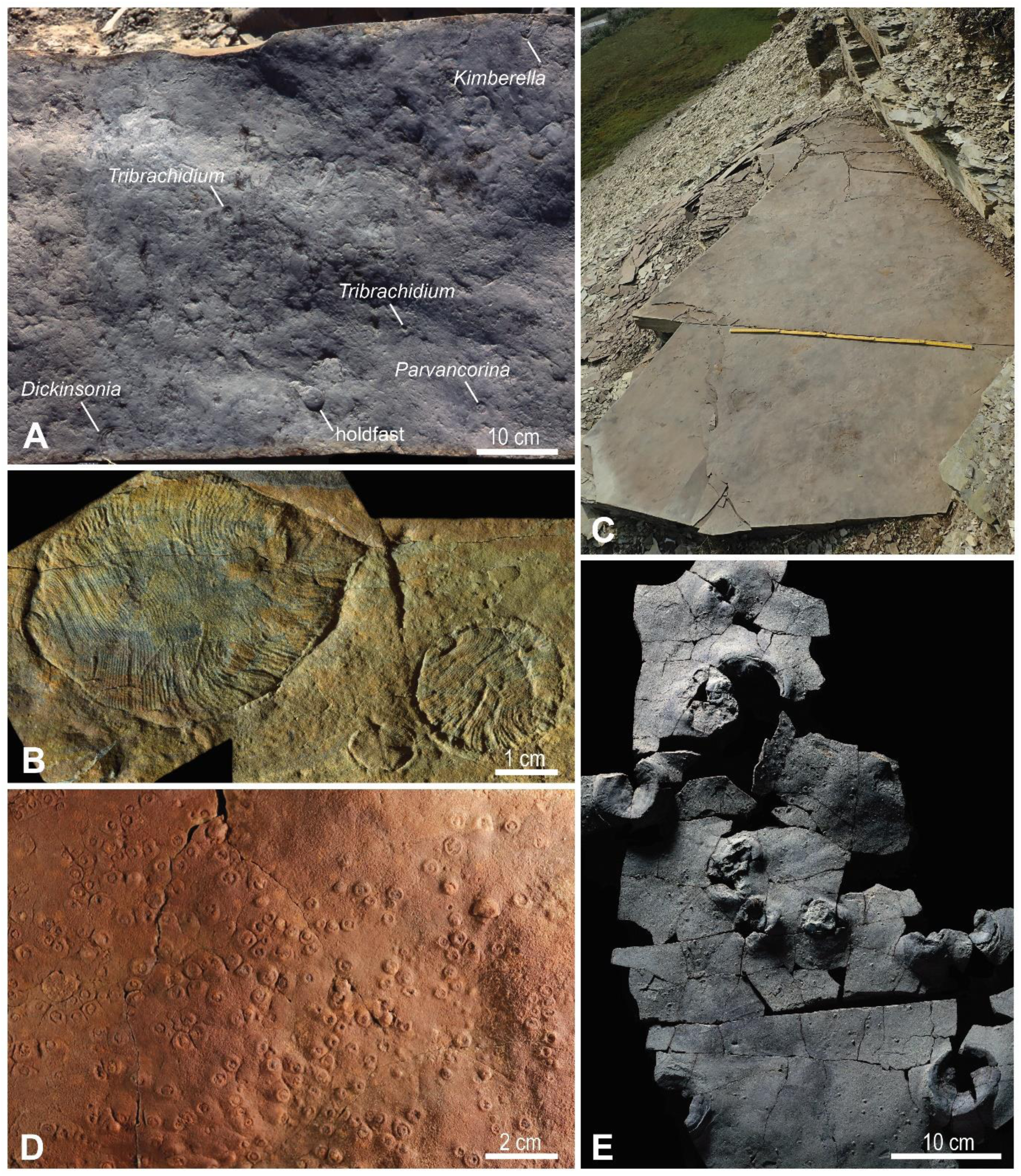
Assemblages of Ediacaran fossils from study localities. A) A fragment of the *Kimberella* surface (KS), indicating key taxa, lower Erga Formation, Winter Coast of the White Sea. B) Specimens of *Dickinsonia* from the *Dickinsonia* surface (DS), Konovalovka Member, Cherny Kamen Formation, Sylvitsa River, Central Urals. C) The *Aspidella* surface (KH1), Khatyspyt Formation, Olenek Uplift, Northern Siberia. Metre rule for scale. D) *Funisia* from FUN4 surface (SAM P55236), Ediacara Member, Rawnsley Quartzite, South Ediacara, Flinders Range, South Australia. E) A representative fragment of the WS surface, upper Lyamtsa Formation, White Sea Region. This particular fragment was not included in the analysis. These data were compared with 7 palaeocommunities that have been subjected to SPPA in previous studies (21, 23), where details of data collection and locality information are described.

**Fig 2.**
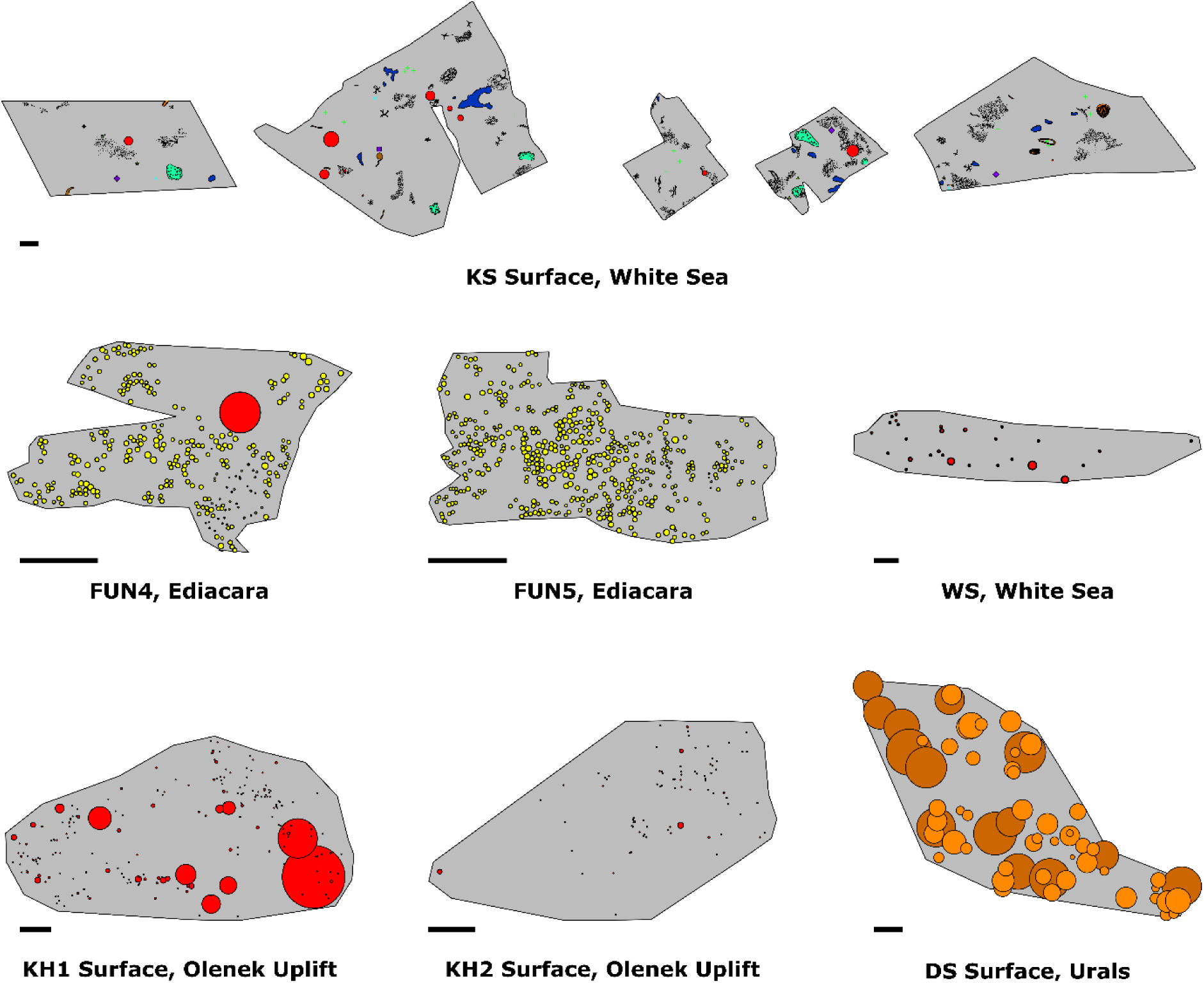
Spatial maps of the seven studied palaeocommunities. Scale bar is 10 cm. Different colours indicate different taxa as follows: Red, *Aspidella;* Orange, *Dickinsonia;* Yellow circles, *Funisia;* Light green scratch marks, *Kimberichnus;* Light green crosses, *Kimberella;* Blue crosses, *Charniodiscus*; Green triangles, *Parvancorina;* Dark blue patches, *Orbisiana;* Black stipples, horizontal traces; White globular strings, *Palaeopasichnus*; Purple diamonds, *Andiva*; Purple squares, *Yorgia*. Size of the circles corresponds to specimen length or diameter (as appropriate). On the DS surface, dark orange circles are the large size-class of *Dickinsonia,* and the light orange represents the small size-class.

**Fig. 3.**
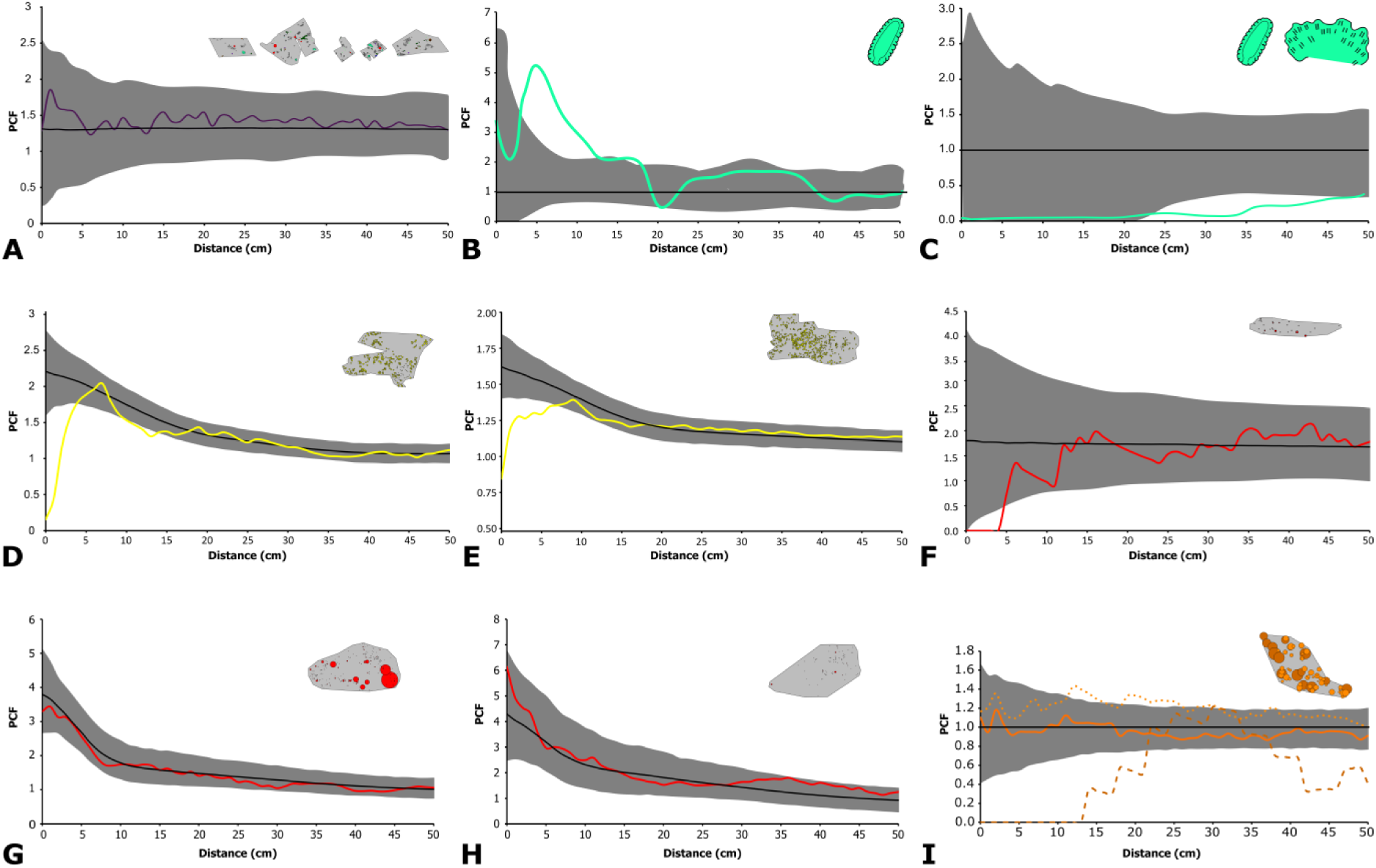
Pair correlation functions describing the spatial distributions of the seven studied palaeocommunities. The coloured lines are the observed data and black lines represent best-fit models (either CSR or heterogeneous Poisson). The grey area is the simulation envelope for 999 Monte Carlo simulations. The x-axis is the inter-point distance between organisms in centimetres. On the y-axis, PCF = 1 indicates complete spatial randomness (CSR), < 1 indicates segregation, and > 1 indicates aggregation. A) The KS surfaces showing sessile specimens with the black-line showing the best-fit heterogeneous Poisson model. B) KS univariate *Kimberella*. C) KS bivariate *Kimberella* – *Kimberichnus* with the CSR model shown. D) FUN4, and E) FUN5 surfaces showing the *Funisia* distributions with the best-fit heterogeneous Poisson model. *Aspidella* from F) WS, G) KH1 and H) KH2 surfaces with their best-fit heterogeneous Poisson models. I) *Dickinsonia* from DS with the solid line showing the whole population, dotted line the juveniles and dashed line the adults with the CSR model shown.

**Table 1.**
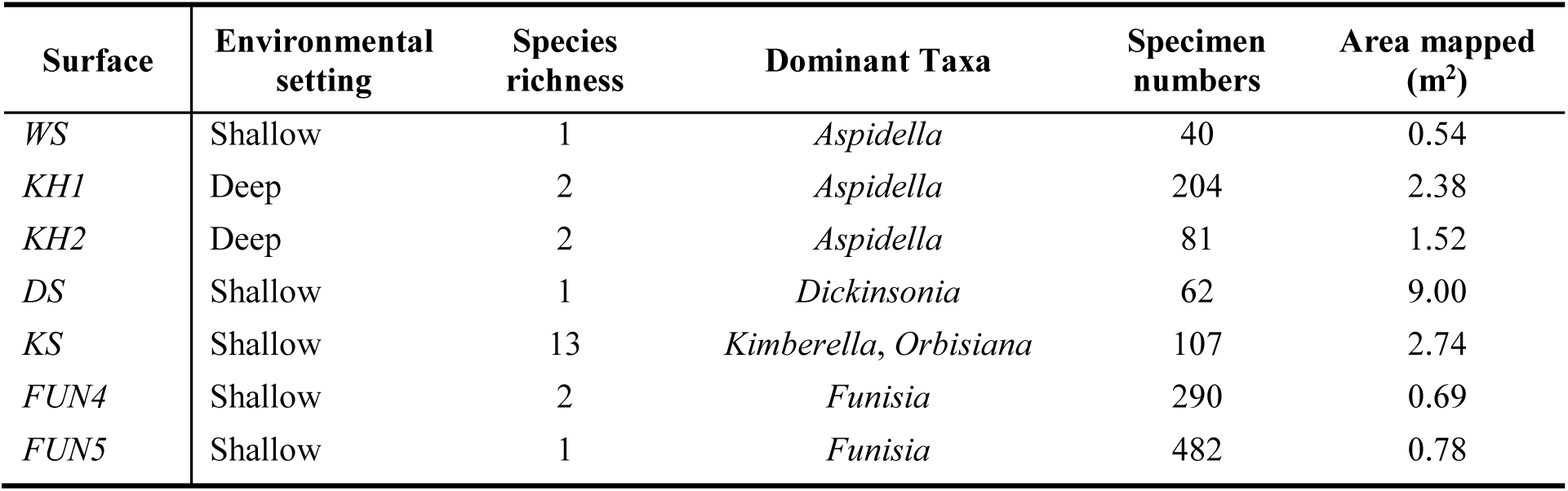
Summary data of the surfaces mapped. The environmental setting, species richness, specimen numbers within the mapped area, and the total mapped area are provided.

**Table 2.** Goodness-of-fit tests for spatial analyses. For the inhomogeneous point processes (HP and ITC), the moving window radius is 0.5 m, using the same taxon density as the taxon being modelled*. p_d_ = 1* corresponds to a perfect fit of the model to the data, while *p_d_ = 0* corresponds to no fit. Where observed data did not fall outside CSR Monte-Carlo simulation envelopes, no further analyses were performed, which is indicated by NA. CSR: Complete spatial randomness indicates, HP: Heterogeneous Poisson model, TC: Thomas cluster model, DTC: double Thomas Cluster, and ITC: inhomogeneous Thomas cluster model. N is the number of specimens mapped. Note that for the mobile taxa *Dickinsonia* and *Kimberella*, and presumed trace fossils formed by mobile taxa (*Kimberichnus*), the observed spatial pattern will also be defined by their behaviour, and so the inference of process from pattern is not as straightforward (see discussion in the main text). The p_d_-value of the best-fit model is given in bold.

| SURFACE | TAXON | N | *P<sub>d</sub>* VALUES PCF |  |  |  |  |
| --- | --- | --- | --- | --- | --- | --- | --- |
|  |  |  | CSR | HP | TC | DTC | ITC |
| WS | *Aspidella* | 40 | 0.019 | **0.796** | 0.504 | 0.2759 | 0.425 |
| KH1 | *Aspidella* | 204 | 0.001 | 0.001 | 0.648 | **0.883** | 0.313 |
| KH2 | *Aspidella* | 81 | 0.001 | 0.001 | 0.576 | **0.932** | 0.001 |
| FUN4 | *Funisia* | 290 | 0.001 | **0.9570** | 0.6340 | NA | 0.245 |
| FUN5 | *Funisia* | 482 | 0.001 | **0.9080** | 0.1320 | NA | 0.218 |
| DS | *Dickinsonia* | 62 | **0.857** | 0.022 | 0.025 | NA | 0.019 |
|  | *Dickinsonia* Small | 48 | 0.128 | **0.978** | 0.143 | NA | 0.158 |
|  | *Dickinsonia* Large | 14 | 0.388 | 0.446 | 0.409 | NA | 0.434 |
| KS | All | 107 | **0.858** | 0.381 | 0.328 | NA | 0.380 |
|  | All sessile | 44 | 0.033 | **0.956** | 0.770 | NA | 0.761 |
|  | *Kimberella* | 18 | 0.001 | **0.837** | 0.491 | NA | 0.103 |
|  | *Orbisiana* | 16 | 0.325 | **0.332** | 0.326 | NA | 0.288 |
|  | *Kimberichnus* | 6 | **0.566** | NA | NA | NA | NA |
|  | Bivariate *Kimberella* – *Kimberichnus* | 24 | **0.028** | NA | NA | NA | NA |

The KS community is notably different in species composition from deep-water communities because it contains mobile organisms such as *Kimberella* and *Yorgia* (96–99) as well as putative trace fossils such as *Radulichus* (thought to be produced by the grazing activity of *Kimberella* specimens) (100). We found that the KS community exhibits CSR, which suggests that any taxon-specific univariate distributions are likely to be biological/ecological in origin, rather than resulting from a taphonomic bias (*p_d_ ^KS^ _All_ =0.858*, Table 2, (23)). In contrast, when all the sessile taxa were grouped together they exhibited a significant aggregation (Table 2), which was best-modelled by a heterogeneous Poisson process (*p_d_ ^KS^ Sessile =0.956,* Table 2). *Kimberella* exhibits a significant aggregation under spatial scales of 20 cm (*p_d_ ^KS^ Kimberella =0.001* for CSR model, Fig. 3A), with Thomas cluster and heterogeneous Poisson models fitting the data well, suggesting that behaviour factors may also influence *Kimberella* spatial patterns. The *Kimberichnus* PCF spatial distribution has a CSR distribution (Fig. 3B, *p_d_ ^KS^ Rad =0.566*, Table 2). Furthermore, the bivariate analyses between *Kimberella* and *Kimberichnus* show a significant segregation (*p_d_ ^KS^ KimRad =0.028*, Fig 3C), which could reflect the *Kimberella* organisms avoiding patches of the surface that had already been grazed.

The *Dickinsonia* population from DS exhibited a CSR PCF distribution (Fig 3I, *p_d_* = 0.857). Analysis of the population of *Dickinsonia* from DS showed two cohorts in the size-distribution (95). The two cohorts exhibited different PCF spatial behavior, with the small specimens aggregating with a best-fit heterogeneous Poisson model (Fig 3I, *p_d_^small^* = 0.978) and the large specimens exhibiting segregation (Fig. 3I).

### Interpreting the spatial distributions of mobile organisms

For mobile organisms, inferring the underlying process behind the observed spatial distributions is imprecise, since their spatial patterns also incorporate contributions from their behavior. Modern animals move primarily to find resources, mates, microhabitats and/or escape predators or detrimental environmental conditions. There is no evidence for predators until the terminal Ediacaran (101), and although we cannot definitely rule out reproductive aggregations, they are also considered unlikely because the largest size-class in the studied *Dickinsonia* population exhibits univariate segregation, so at time-of-burial, the organisms were not aggregating as might be expected in a mating event. Furthermore, the majority of extant marine benthic organisms use broadcast spawning to reproduce sexually (102), so do not require the two mating organisms to be within the spatial scale (< 40 cm) found on the DS surface. We cannot determine whether the large *Dickinsonia* are reacting to the mortality event which killed and preserved them, however, this would not explain the complex interplay between aggregation and segregated behaviors. Therefore, for this *Dickinsonia* population, the search for resources and/or microhabitats is considered most plausible explanation, particularly since this hypothesis is further supported by their spatial patterns. Aggregated – segregated PCF patterns such as those seen in our *Dickinsonia* population are common in extant sessile organisms where juveniles are initially aggregated on preferred habitats but then begin to compete with each other as they require greater resources, leading to thinning or segregation amongst adult populations (55). While it is not possible to confirm the underlying mechanism for the distribution of the studied *Dickinsonia* population, we consider it most likely to be motivated by associations with preferential habitat for food and/or resources. Further analyses of other *Dickinsonia* surfaces would enable more robust conclusions to be reached.

### Time averaging

The preservation of time-averaged communities has the potential to bias our analyses (see (21, 25). In Avalonian communities, taphomorphs interpreted to record the decaying remains of organisms are identified by their poor preservational fidelity, irregular morphologies, and often high topographic relief (103). This interpretation is consistent with data suggesting that the spatial interactions of some taphomorph populations mirror those of other taxa they are considered to be derived from (21). Taphomorphs are considered unlikely to have imparted a significant signal on these studied surfaces, since we did not observe ivesheadiomorph-type forms, and there is a consistent level of preservational detail amongst fossil communities

*Funisia* communities tend to have very similar diameters for the holdfasts, which suggests single colonization events (104). Different reproductive events can be distinguished by population analyses of size-distributions (105), with each reproductive event identified through statistically significant cohorts within the size-distribution (90). Surfaces FUN4 and FUN5 both exhibit populations with two cohorts (SI Figure 1), most likely indicating two reproductive/colonization events. The best-fit models for each of these surfaces are heterogeneous Poisson models (Fig. 3, Table 2), with very high goodness-of-fit values (*p_d_ > 0.90*) reflecting a single model for each surface. Therefore, cohorts of *Funisia* specimens on each of the studied surfaces were affected by the same underlying environmental heterogeneity, so most likely were contemporaneous.

## Discussion

The univariate and bivariate analyses of five out of seven of the studied palaeocommunities provide compelling evidence that their local environment had a significant influence on their communities (Fig. 3, Table 2). In modern settings, habitat associations form when a patchy resource provides heterogeneously distributed preferential conditions for the establishment and growth of sessile taxa, and/or feeding ‘hotspots’ for the mobile taxa (47,54,83). The presence of inferred habitat interactions within our palaeocommunities showed a significant correlation with the environmental setting (Kruskal-Wallis Test, *p = 0.049*), with all five palaeocommunities with strong habitat interactions derived from shallow-water settings. The two communities that were seemingly not strongly influenced by their local habitat are from deep-water facies (Table 2). These results are consistent with previous work, which found that for seven independent deep-marine (slope and basin) Ediacaran palaeocommunities from Newfoundland and Charnwood Forest, only one was dominated by associations of taxa with local habitat heterogeneities (21–23) (Kruskal-Wallis Test of all data, *p = 0.021*; Fig. 4).

**Fig. 4.**
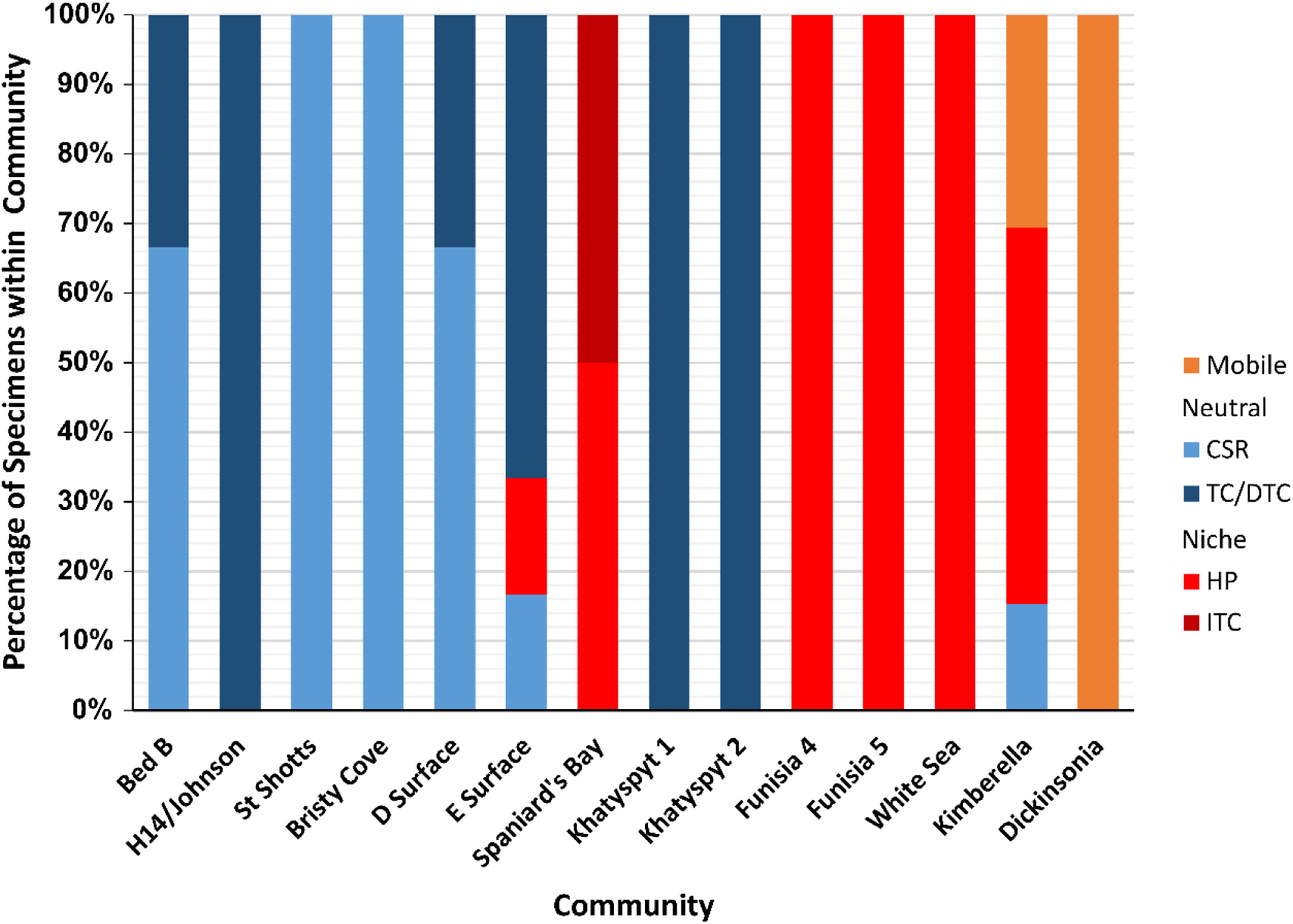
Proportion of best-fit univariate models by surface, adapted from (23). The percentage of specimens within the community with univariate spatial distributions that are best described by CSR, HP, TC (or DTC) and ITC models. CSR and TC are considered random or dispersal (neutral) models and are shown in blue. HP and ITC are local environmentally driven (niche) models, shown in red. Mobile taxa are shown in orange, and inferred to be environmentally-driven. Data and plot for surfaces Bed B to Spaniard’s Bay from ref. (23).

**Fig. 5.**
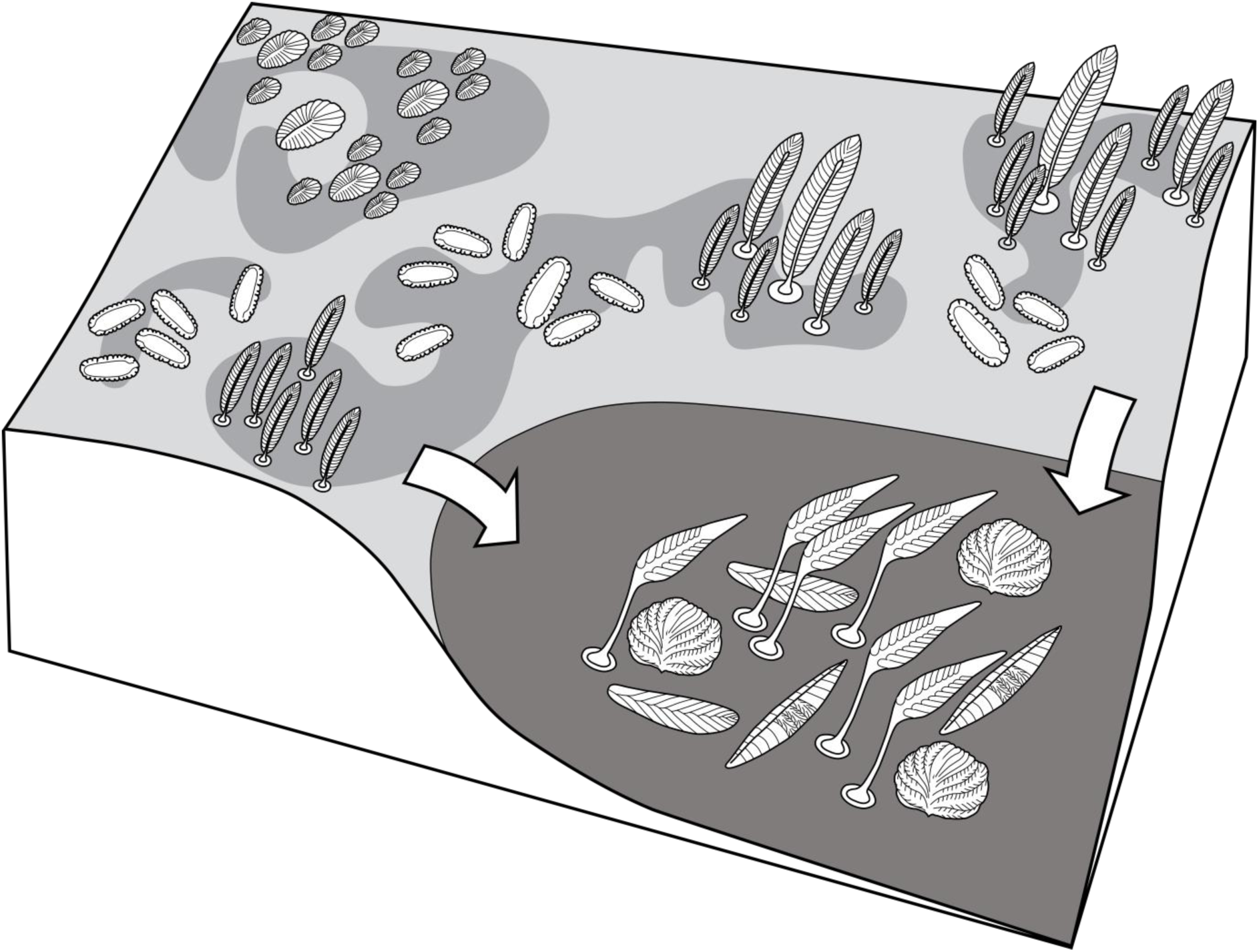
Schematic diagram showing variation of heterogeneities within different environmental settings. Shallow water communities are significantly influenced by habitat heterogeneities. Grazing within these shallow waters further increases substrate heterogeneity, potentially increasing diversification. Furthermore, this grazing increases deep-water heterogeneity through the creation of different sized particulate organic matter due to the influx of particulate matter from the shallows.

Untangling environmental from evolutionary trends in the Ediacaran has been hampered by a limited overlap between temporal periods and environmental settings (1, 17). The palaeocommunities in this study derive from successions within a variety of lithologies (tuff, coarse sandstone, mixed siltstone, limestone) as well as palaeogeographic positions (17,62,63,69,104,106,107). We find no significant direct correlations between these factors and the relative importance of habitat heterogeneities on the studied surfaces (*p >> 0.1;* Fig. 3, Table 2). The palaeocommunities that are not influenced by local habitat heterogeneities (KH1 and KH2) are hosted within carbonate successions (107), making them distinct from the siliciclastically-hosted palaeocommunities on the KS, WS, FUN4, FUN5 and DS surfaces, or in previous (21–23) work. However, the Khatyspyt surfaces behave ecologically in the same way to Avalonian palaeocommunities derived from similar depths, but different lithological successions (21–23), suggesting that lithology alone is not causing the KH1 and KH2 surfaces differing results. Therefore, two possible factors remain that may explain the differences in community dynamics found here. The differences could reflect evolutionary trends, and it is true that the oldest studied palaeocommunities show limited habitat influence (21–23), when compared to the younger palaeocommunities documented in this study (Fig. 4). Unfortunately, the lack of fine-scale dating across these communities and older Avalonian ones precludes detailed fine-scale regression to assess whether either the Khatyspyt palaeocommunities are an outlier to this apparent trend, or this trend merely reflects the biases of the available data. Alternatively, the differences could be due to the environmental setting. We have shown that Ediacaran environmental setting has a significant influence on community dynamics (*p = 0.021*), with shallow water palaeocommunities significantly influenced by habitat heterogeneities, in contrast to the deep water palaeocommunities (Fig 3, Table 2; (21–23)).

While SPPA have only been applied to a small proportion of the known in-situ Ediacaran palaeocommunities (17 studied surfaces (21–23, 23, 60, 108)), there is a notable correspondence between the importance of habitat heterogeneities to community ecology and assemblage diversity. In this study, the palaeocommunities exhibiting significant influence from local habitat heterogeneities are those that belong to the diverse White Sea assemblage, which is in contrast to the previous work on Avalonian palaeocommunities (21–23), which are not significantly influenced by such heterogeneities. The relationship between environmental spatial heterogeneities and species richness is well established, with habitat variations enabling species co-existence through the creation of different niches (109). This relationship extends to modern deep-sea benthic communities, where these heterogeneities have been shown to provide a mechanism for diversification on large scales, such as between canyons, trenches, seamounts (110, 111), on the centimetre to metre scale (112), and through microhabitats (45).

Tentatively, we propose that the ecological differentiation observed between Ediacaran shallow and deep-water communities may evidence the late Ediacaran development of a chain of evolutionary diversification. This chain started in shallow water communities, with the creation of habitat patchiness by mobile Ediacaran organisms, which then led to a feedback of increasing diversification that ultimately expanded into the deep-sea. This hypothesized feedback could have promoted diversification throughout the Ediacaran by increasing heterogeneity as follows:

First, metazoan mat grazing creates spatial heterogeneity in microbial substrates through the formation of depleted and non-depleted patches (113). Our data suggest that once created, organisms such as *Kimberella* may have avoided pre-grazed patches, with this selective grazing accelerating further creation of mat heterogeneity (Fig 3C). Secondly, the grazing-induced creation of different-sized detrital particles in the form of differential-sized fecal pellets and fragments of non-consumed food within the water-column (114), would have created new food sources and therefore potential new niches. Thirdly, this shallow-water differentiated particulate organic carbon (POC) and matter (POM) could have eventually filtered through to deep-sea communities, promoting deep-sea heterogeneity. In the modern ocean, the main source of deep-sea habitat heterogeneity is small-scale variation due to differentiated particle influx (114), with the majority of the particulate organic carbon (POC) coming from phytodetritus, which is transported from shallow waters to deep waters by ocean currents, tides and upwelling (114, 116). In the modern ocean, the diurnal vertical migration of mesozooplankton and macrofauna contributes up to 50% of POC to the deep-sea via fecal pellets (116–118). A planktonic/larval stage for Ediacaran organisms has been predicted on the basis of their likely waterborne dispersal mechanisms (25, 105), but there is presently no direct evidence of non-larval, planktotrophic zooplankton until the onset of the Cambrian (119). In the absence of planktotrophic zooplankton and macrofauna, the Ediacaran POC flux may have been either larger, due to lack of consumption of phytoplankton in the shallow water, or smaller, due to a lack of mixing by diurnal vertical migration of the plankton (6), and this cannot yet be determined. However, the other ∼50% of POC flux in the modern oceans is transported from shallow to deep-water via oceanic currents and upwelling (114, 116), which should still have operated in the Ediacaran. However, prior to grazers and detritivores, this POC/POM flux would have been relatively homogenous phytodetritus. The evolution of grazers would have led to a shift towards size differentiated POC/POM, potentially increasing the heterogeneity of the deep-sea landscape (114), and providing a mechanism for deep-marine diversification.

Budd and Jensen (12) introduced the Savannah hypothesis to explain early animal diversification, whereby Ediacaran diversification was driven by small-scale variations in local habitat. They argued that it was the drive to find these heterogeneous distributed resources that led to novel evolutionary innovations such as mobility. Our results demonstrate that at least some of these early animal communities that contain mobile organisms were influenced by such habitat variations, and we describe a mechanism that links early animal diversification and benthic habitat patchiness prior to the evolution of predators and wide-spread pelagic organisms. We show that taxa such as *Kimberella* had a segregated distribution with trace fossils considered to be their grazing traces (98), suggesting that they may have been capable of avoiding non-preferred areas, possibly already consumed patches, revealing adaptation of behavior when interacting with these patches. This adaptation theoretically has the capacity to drive further diversification, initially dependent on the environmental-setting, starting in the shallow water, and then, over time, moving into deeper water, but currently available global fossil assemblages limit the testing of this prediction. If this hypothesis is correct, we would expect deep-water assemblages to diversify during the terminal Ediacaran and into the Cambrian. Our results therefore provide tentative support for the Savannah hypothesis, suggesting that this late Ediacaran taxonomic diversification was a benthic event, which facilitated a chain of diversification by promoting marine habitat heterogeneities.

## Conclusions

We present evidence to suggest that the influence of local habitat on Ediacaran organisms is significantly correlated with broad-scale environmental setting. The relationship of Ediacaran communities to habitat-dependent interactions is correlated with Ediacaran assemblage diversity, with communities from the more diverse White Sea assemblage showing significant habitat associations and interactions in contrast to relatively habitat insensitive deep-sea Avalonian assemblages. We suggest that the presence of shallow-water grazers could have created further habitat heterogeneity in shallow-water and ultimately deep-water, via the heterogenization of the shallow-water substrate and via the introduction of variable size particulate matter to the deep-sea. These results demonstrate the utility of these approaches for investigating the early diversification of metazoans. We have shown the importance of local environmental patchiness to the diversification of early animals, and our results are consistent with the hypothesis that the early diversification of metazoans was a benthic event, driven by responses to habitat patchiness.

## Acknowledgements

We thank K. Nagovitsin and O. Zharasbayev (IPGG SB RAS) for help with mapping surfaces KH1 and KH2, and J. Gehling and M. Binnie of the South Australia Museum for assisting with access to Australian material.

## Funding

This work has been supported by the Natural Environment Research Council [grant numbers NE/P002412/1 and Independent Research Fellowship NE/S014756/1 EGM, and Independent Research Fellowship NE/L011409/2 to AGL], a Gibbs Travelling Fellowship (2016-2017) from Newnham College, Cambridge, and a Henslow Research Fellowship from Cambridge Philosophical Society to EGM (2016-2019). Field research in the White Sea Region, Arctic Siberia and Central Urals has been supported by the Russian Science Foundation [grant number 17-17-01241 to DG]. SX acknowledges funding from the NASA Exobiology and Evolutionary Biology Program [80NSSC18K1086]. Large image processing and interpretation of photomontages of the *Dickinsonia* Surface was supported by the Russian Foundation for Basic Research [grant number 19-05-00828 to AVK].

