## Supplementary Figure 1 for "The influence of environmental setting on the community ecology of Ediacaran organisms"

### Supplementary Material

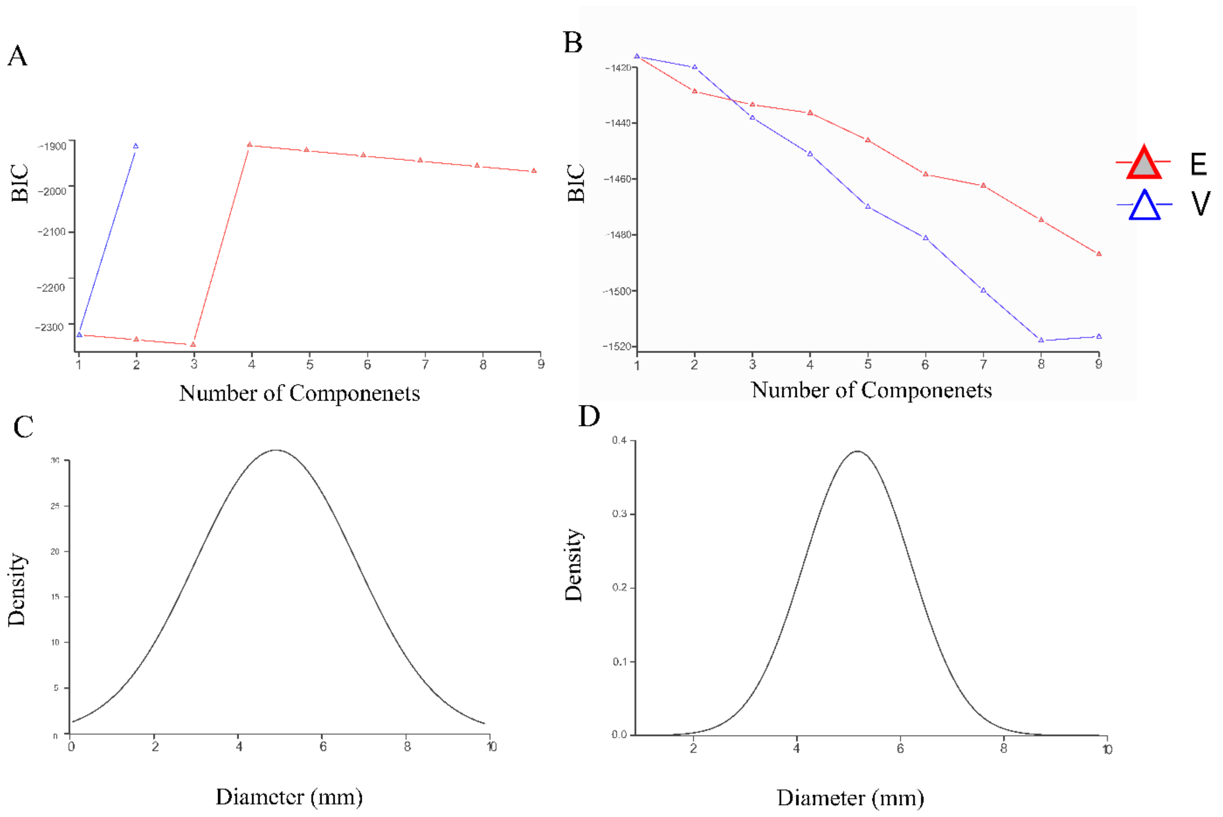


Figure S1: Analyses of the size-distribution of the *Funisia* populations from FUN4 and FUN5. BIC values for A) FUN4 and B) FUN5 showing strongly significant (>10) two-cohort best-fit models for both surfaces. Density distributions for C) FUN4 and D) FUN5 surfaces, demonstrating that the second cohorts represent a small proportion of the total populations.
